# Microfluidic Control of Dorsal-Ventral Patterning Within a Single Forebrain Organoid

**DOI:** 10.64898/2026.04.05.716567

**Authors:** Sebastian Torres-Montoya, Samira Vera-Choqqueccota, Spencer T. Seiler, David Haussler, Sofie R. Salama, Mohammed A. Mostajo-Radji, Mircea Teodorescu

## Abstract

How distinct regional identities emerge within a single developing brain remains poorly understood. Current *in vitro* models address this by fusing independently generated organoids, but this introduces variability in size, maturation state, and connectivity, confounding the study of regionalization itself. Here, we present a microfluidic platform that supports the co-development of different tissue identities within a single, continuous 3D culture domain. The device integrates controlled microfluidic flow with real-time fluorescence imaging, providing stable perfusion and high-resolution tracking of molecular transport without the need for embedded sensors or disruptive sampling. By delivering SAG, a Sonic hedgehog pathway agonist, to one surface of mouse forebrain organoids, we induced spatially segregated ventral (Nkx2.1^+^) and dorsal (Pax6^+^) domains within a unified tissue architecture. Controlled morphogen delivery is sufficient to drive region-specific fate specification without organoid fusion, offering a practical, scalable alternative for studying tissue regionalization *in vitro*.

## INTRODUCTION

Recent advances in 3D cell culture technologies have enabled the creation of *in vitro* models that closely reflect tissue microenvironments^1^, improving disease modeling^2^ and drug-testing platforms^3,4^. Despite progress in organoid models, achieving controlled regionalization within a single organoid, particularly in the brain, remains a significant challenge. Most current strategies rely on generating multiple organoids with distinct cellular identities and fusing them into “assembloids” ^5,6^. This process introduces variability in maturation, geometry, and interaction dynamics, limiting reproducibility and experimental control.

A significant problem for regionalization studies is the limited ability to precisely control and measure low-concentration biomolecules, such as nutrients^7^, metabolites^8,9^, and gases^10,11,12^, inside 3D tissues^13^. These molecules drive patterning, boundary formation, and metabolic specialization. Biomolecule distribution is difficult to quantify in real time, and spatial resolution is difficult to predict; both are essential for studying regional identity formation. Some biosensing approaches, such as impedance-based systems^14,15,16^, optochemical oxygen sensors^10,12^, and integrated electrochemical detectors^8,9,17^, have expanded *in situ* analysis^18,19^, but they typically provide bulk or endpoint measurements and often require complex hardware integration. Platforms designed for monitoring oxygen^11,20^, glucose^9^, or neurotransmitters^21^ rely on invasive electrode arrays that are more difficult to adapt across tissue models^22^. While organ-on-chip platforms^8,17,19,23,24,25,26^ and advanced imaging workflows^27,28^ have improved access to microenvironmental analysis, few tools enable non-destructive, high-resolution mapping of low-concentration gradients within heterogeneous 3D tissues^27,29^. Such capability is required to precisely measure the regionalization process in one organoid. Fluorescence microscopy has the required sensitivity, but its use for analyte mapping in 3D microfluidic systems remains limited^8,17,23,30^.

To address these challenges, this work introduces a microfluidic platform specifically designed to establish and study regionalization within a single organoid. The device uses controlled perfusion and continuous fluorescence imaging to modulate and visualize low-concentration molecular environments without embedding sensors or disturbing the tissue. The vertical-chamber geometry ensures stable culture positioning and full optical access, enabling precise, real-time mapping of how diffusible components shape local microenvironments. By controlling localized molecular cues within a single continuous 3D construct, this platform creates the conditions necessary for distinct regional identities to emerge within a single organoid, offering a direct alternative to assembloids. Instead of fusing separately generated organoids, we can now guide region-specific development using morphogens within a single unified tissue architecture. This approach represents a shift from assembling organoids toward actively growing integrated organoid architectures^5,6^, thereby opening new opportunities for studying spatial patterning, metabolic specialization, and early brain regionalization *in vitro*.

## RESULTS

### Platform description

The microfluidic imaging platform was developed as a modular, compact device to enable dynamic observation and environmental control of biological samples^25^. The configuration enables standalone operation in experimental workflows that require live-cell manipulation. The fluidic subsystem utilizes a PDMS/glass microfluidic chip (Fig. 1a-c) that drives flow through a syringe pump; see description methods. The platform integrates multiple subsystems, including fluid handling, environmental regulation, and real-time fluorescence imaging, each designed for integration on an optical breadboard (Fig. 1d). This arrangement enables precise control of fluidic conditions around the sample while maintaining consistent flow. The chip is in contact with a thermal control that includes a heater pad and an embedded temperature sensor (Fig. 1f), allowing temperature regulation across the chip. The chip’s inlet and outlet ports facilitate directed flow (Fig. 1g) while enabling video recording.

**Figure 1.**
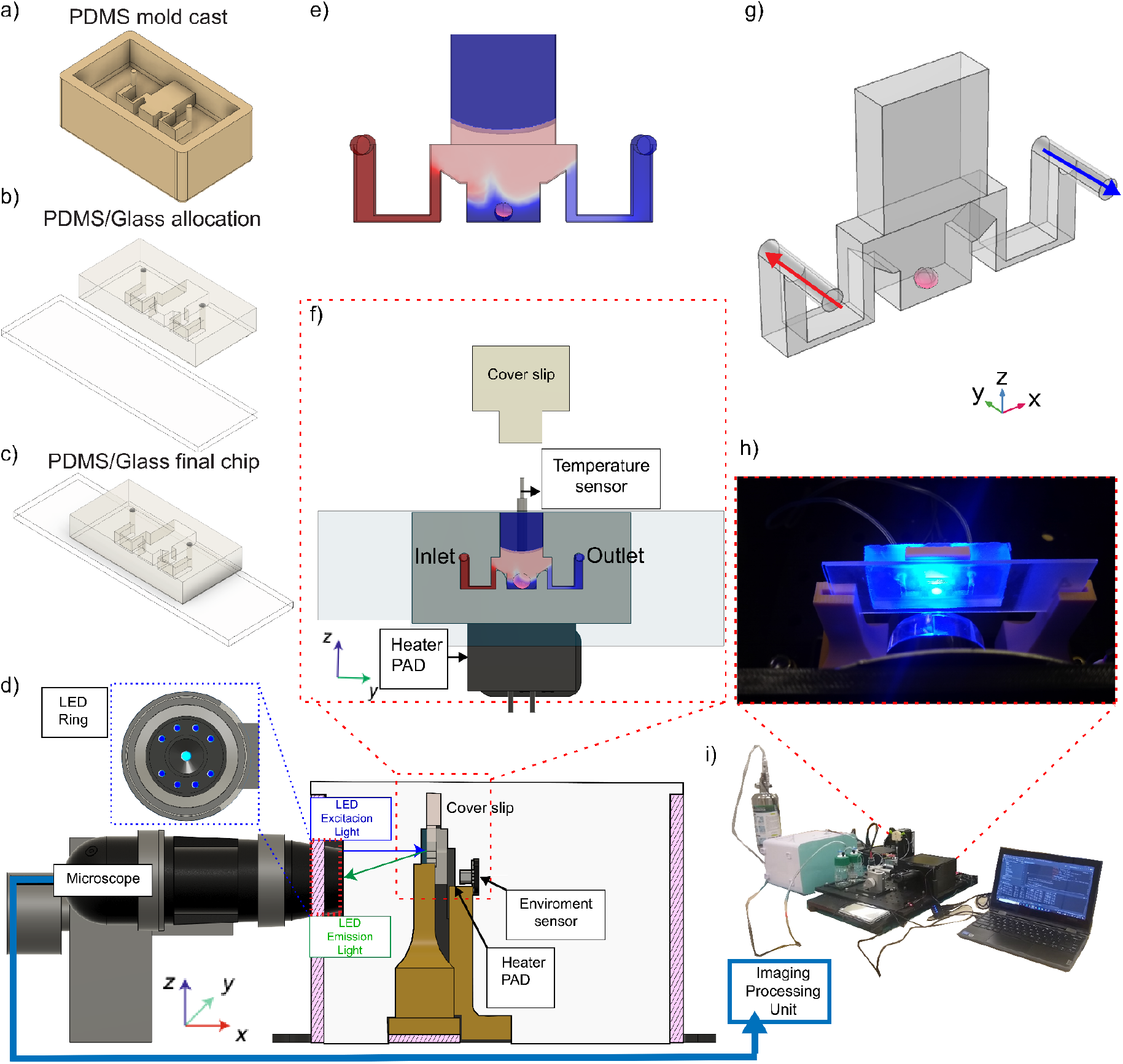
Schematic and experimental setup of an integrated microfluidic fluorescence imaging platform. (a) PDMS mold used to generate the microchannel geometry. (b) Assembly of the molded PDMS structure onto a glass substrate. (c) Final bonded PDMS/glass chip after alignment and sealing. (d) Schematic of the assembled platform, (e) 3D simulation rendering of the concentration distribution within the chip. (f) 3D rendering of the chip showing the flow distribution, (g) fluid flow directions of the experiments. (h) An optical detection module composed of a microscope, an LED excitation ring, and an emission light path. (i) Fully assembled laboratory setup with the imaging processing unit, electronics, and peripheral control subsystems. Supplementary Fig. 5 provides a detailed description of the platform.

The imaging module (Fig. 1h) comprises a fluorescence-capable microscope, including an LED ring light, dichroic and emission filters, and a high-resolution camera connected to a processing unit. The blue excitation and green emission light are clearly defined and adapted for minimal signal loss. Environmental sensors are located within the enclosure to monitor pH, temperature, and fluid flow, enabling responsive adaptation to changing experimental conditions. Within the PDMS microchannel structure (Fig. 1f,h), objects of interest are illuminated under a microscope. This configuration enables live fluorescence imaging while maintaining stable flow and minimal perturbation. The platform enables optical observation and sample placement, ensuring spatial and temporal resolution.

The physical configuration of the platform presents the illuminated microfluidic chip during operation (Fig. 1h), which includes real-time image display, experimental control software, and the hardware arrangement incorporating the fluidic and optical subsystems (Fig. 1i). The entire platform was validated through continuous imaging trials, confirming reliable excitation-emission detection. These results establish the platform’s applicability for the record of biological assays that require integrated environmental and optical control.

### Computational Fluid Dynamics (CFD) study of the spatial distribution of low-concentration molecules

To evaluate the microfluidic platform’s ability to generate spatial concentration gradients, we conducted a detailed spatiotemporal simulation of solute concentration under pulsatile flow conditions. A pulsatile delivery protocol was simulated using Finite Element Analysis (FEA) software, with a green fluorescent dye at physiologically relevant concentrations. The simulation replicated the injection of an analyte (50 nM) under laminar-flow conditions, every 300s, see Fig.1g. Simulations were conducted using custom boundary conditions applied to the microfluidic chip geometry and the experimentally observed flow rates, as shown in Supplementary Table 1 and Supplementary Fig. 1.

The simulation illustrates how solute enters from the left inlets. Snapshots taken every 600 seconds from t = 0 to 4000 s show how the concentration, Fig 2a, measured in nanomolar (nM), evolves, from uniform low levels (0 nM) to peak values around 50 nM. An initial concentration of 50 nM was injected at 0.005m/s until the domain reached the final peak concentration. Additionally, we plot the concentration distribution across different cross-sectional planes (Fig. 2g) to detail the compound’s absorption process in the organoid (Fig. 2d-f). Over time, the solute distribution becomes non-uniform, with limited mixing at the center of the chip. A cross-sectional view highlights six key regions of interest (ROIs): Top, Bottom, Left, Right, Middle, and Center. This view shows how solutes accumulate differently across the organoid. The Top and Right regions reach higher concentrations more quickly, whereas the Bottom and Left regions exhibit delayed and lower solute delivery (Fig. 2c).

**Figure 2.**
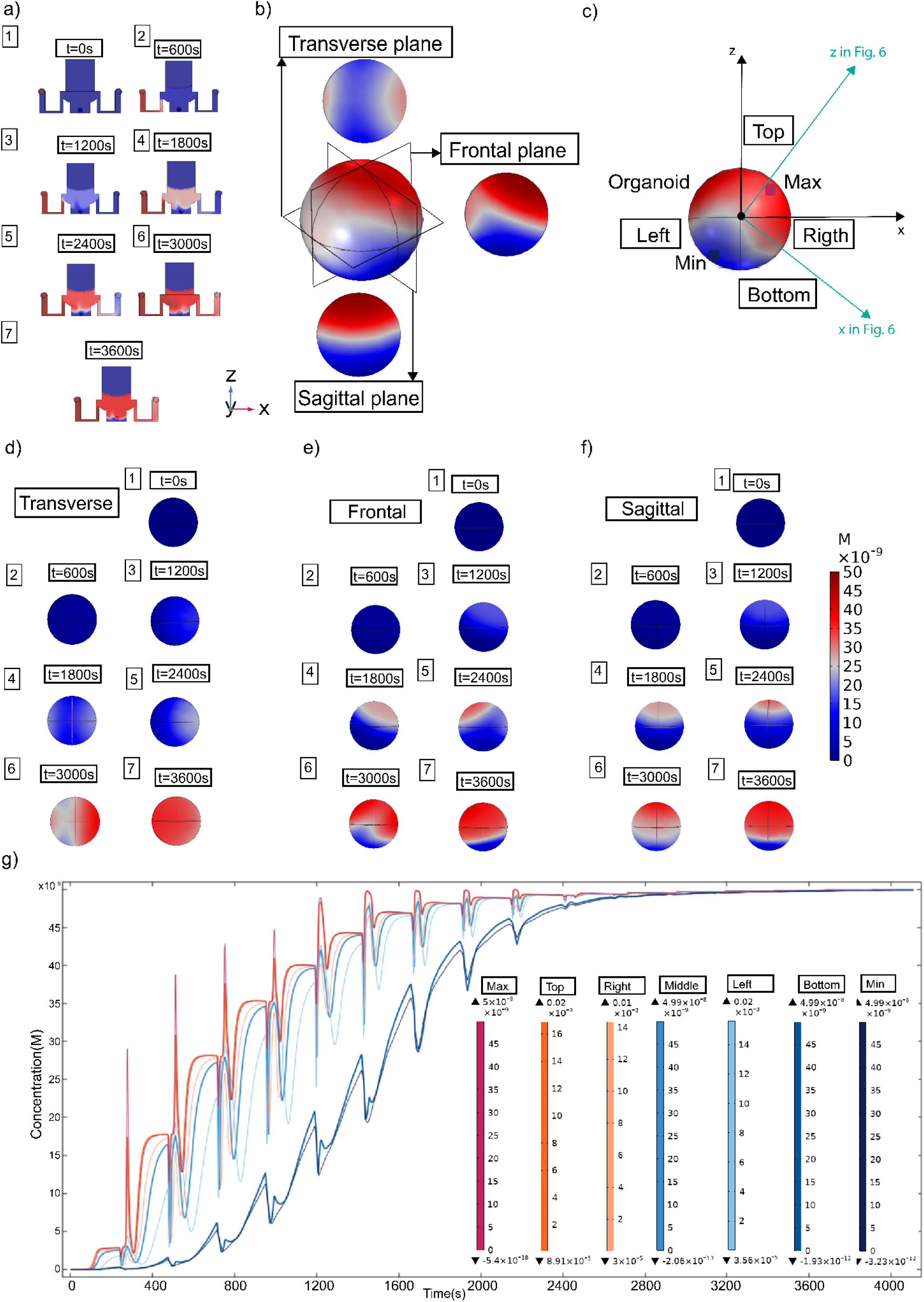
Computational modeling of solute transport dynamics in a pulsatile microfluidic environment across multiple spatial planes. (a) Cross-sectional views of the device at increasing time points (1. 0s, 2. 600s, 3. 1200s, 4. 1800s, 5. 2400s, 6. 3000s, 7. 3600s, showing the progressive entry and distribution of the diffusing species from the inlet (blue) toward the outlet (red). Concentration values in this representation range from 0 to 50 nM, with higher intensities localized near the inlet at early time points. (b) Auxiliary plane views. (c) CFD selected concentration points showing the real experimental allocation of the organoid in the platform (arrows in blue), see Fig. 6. (d) Transverse plane view of the concentration profile at steady state, revealing the formation of a gradient across the chamber cross-section. (e) Frontal plane view of the concentration profile, showing concentration differences along the inlet-outlet axis, spanning 0-50 nM. (d(1-7)) Time-resolved concentration maps in the transverse plane at 0, 600, 1200, 1800, 2400, 3000, and 3600 s, demonstrating how the gradient evolves toward partial homogenization while still maintaining localized differences of several millimolar between top and bottom regions. (e(1-7)) Time-resolved concentration maps in the frontal plane at 0-3600s, showing the gradual establishment of the molecular gradient from inlet to outlet, with concentration increasing steadily in the downstream region. (f(1-7)) Time-resolved concentration maps in the sagittal plane at 0-3600s, providing orthogonal confirmation of diffusion patterns, with concentration values progressing from nanomolar levels at the chamber periphery to millimolar levels near the main channel. Color bar indicating normalized concentration, spanning 0 nM (blue) to 50 nM (red). (e) Absorption concentration profiles derived from the frontal plane (g), showing temporal dynamics during the first 2400s of the simulation. Regional measurements (top, right, middle, left, and bottom surfaces) confirm that concentrations increase from nanomolar at early time points to millimolar as diffusion progresses, with profiles converging toward a stable distribution.

The time-dependent concentration response in six defined regions (Top, Right, Middle, Left, Bottom, and extremes Min/Max) is plotted in the bottom pane, Fig. 2g. Concentration is measured in nanomolar (nM), with peak pulse values reaching ∼45-50 nM in Top and Right, and delayed, lower-amplitude responses (∼15-30 nM) in Left and Bottom. These values are consistent with previous work^25,31^. The middle zone stabilizes at approximately 40 nM. These results confirm the platform’s ability to generate resolved, pulsatile, and asymmetric solute distributions, thereby enabling studies that require localized compound distributions, environmental control, and differential exposure in organoid cultures.

To track concentration changes over time, we plotted time-resolved profiles (nanomolar [nM]) for each ROI over a 4,000s simulation. The Top and Right areas exhibit a compound uptake, pulsatile peaks (40-50 nM), whereas the Bottom and Left areas show slower and weaker responses (15-30 nM). The Middle region has intermediate values. Each ROI shows peak and baseline levels, with maxima ranging from 0.05 nM in high-exposure zones to 4.99×10^−8^ nM in low-exposure zones. Minimum values in some areas drop close to 8 × 10^−8^ nM, indicating a very steep concentration gradient (Fig. 2g). Overall, these results confirm that the platform can produce precise microenvironments. This level of spatial and temporal resolution is helpful for studies needing localized delivery, directional stimulation, or concentration-specific cellular responses.

### Experimental verification of the CFD study using forebrain organoids

To validate the CFD results, we conducted experiments using mouse forebrain organoids in the microfluidic device and introduced a green fluorescent tracer. Dye gradient formation and quantification were then characterized within the device to verify the computational predictions (Fig. 3). The experimental setup is illustrated schematically, including the device design, the BODIPY green fluorescent tracer, three defined regions of interest (ROIs), and the optical detection configuration (Fig. 3a). To validate fluorescence concentration, a calibration curve was generated using 50 nM dye aliquots, which demonstrated a linear relationship between concentration and fluorescence intensity (R^2^ = 0.9423), as shown in Fig.3b, thereby confirming both the robustness and stability of the generated gradients. Quantitative analysis of the concentration profiles in the top, middle, and bottom ROIs confirmed consistent gradient formation, as evidenced by measurements from three independent replicates. The resulting profiles showed reproducible behavior over time Fig.3 (c-e). The time-resolved concentration measurements across ROI 1-3 confirmed the stable formation of a gradient, Fig.3 (c-e). Differences between regions of interest over time, demonstrating spatial concentration offsets that remain stable during the experiment, Fig.3 (f-h). This validation supports the use of the platform for controlled delivery studies under physiologically relevant low-concentration conditions. The calibration values are presented in detail in Supplementary Table 2. Time-lapse fluorescence imaging captured the spatial and temporal evolution of the gradient at different time points (t = 0 min, 42 min, and 83 min), revealing the progressive establishment of a stable dye distribution across the organoid (Fig. 3i-k). Additionally, details of the quantification process are shown in Supplementary Fig. 2.

**Figure 3.**
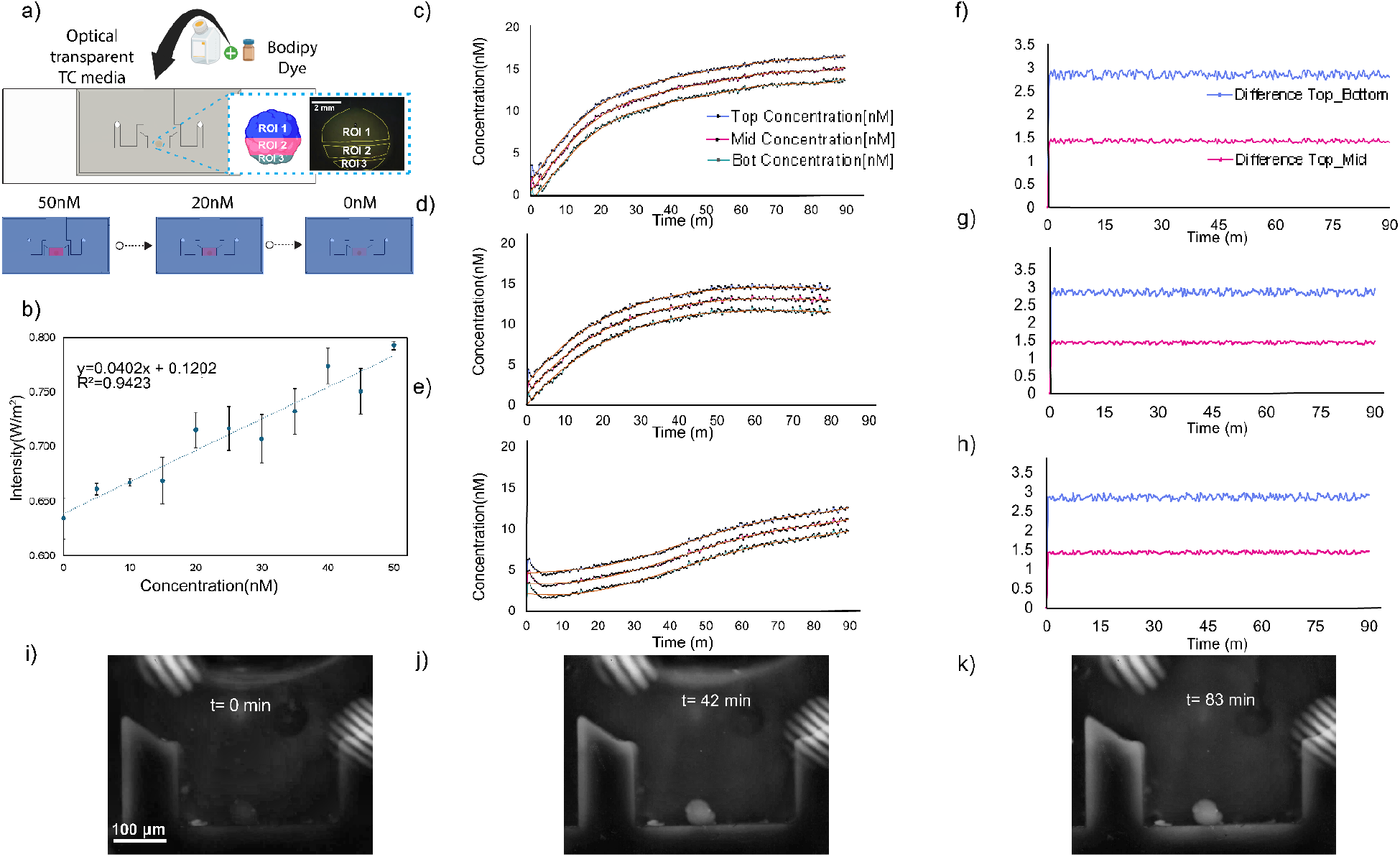
Characterization of BODIPY dye distribution and gradient formation in the microfluidic device. (a) Schematic of the experimental setup showing device design, regions of interest (ROI 1-3), and the optical detection platform, representative simulation snapshots of concentration fields at 50 nM, 20 nM, and 0 nM. (b) Calibration curve relating dye concentration (0-50 nM) to fluorescence intensity, showing a linear correlation (R^2^ = 0.9423). (c-e) Time-resolved concentration measurements across ROI 1-3 at an initial concentration of 50nM, confirming the stable formation of a gradient. (f-h) Differences between regions of interest over time, demonstrating spatial concentration offsets that remain stable during the experiment. (i-k) Fluorescence microscopy images of an organoid within the device at t = 0 min, 42 min, and 83 min, confirming dye distribution and gradient persistence.

### Case study: cell migration and organoid regionalization using morphogens

To assess the platform in a physiologically relevant use case, we cultured mouse forebrain organoids on the microfluidic device starting at day 3, following neuronal induction^31^. Organoids were exposed for two days to SAG, a Sonic hedgehog (Shh) pathway agonist known to promote ventral forebrain identity^32^, and analyzed at day 5. Immunofluorescence analysis revealed robust neuronal differentiation and spatial regionalization within individual organoids (Fig. 4c-h).

**Figure 4.**
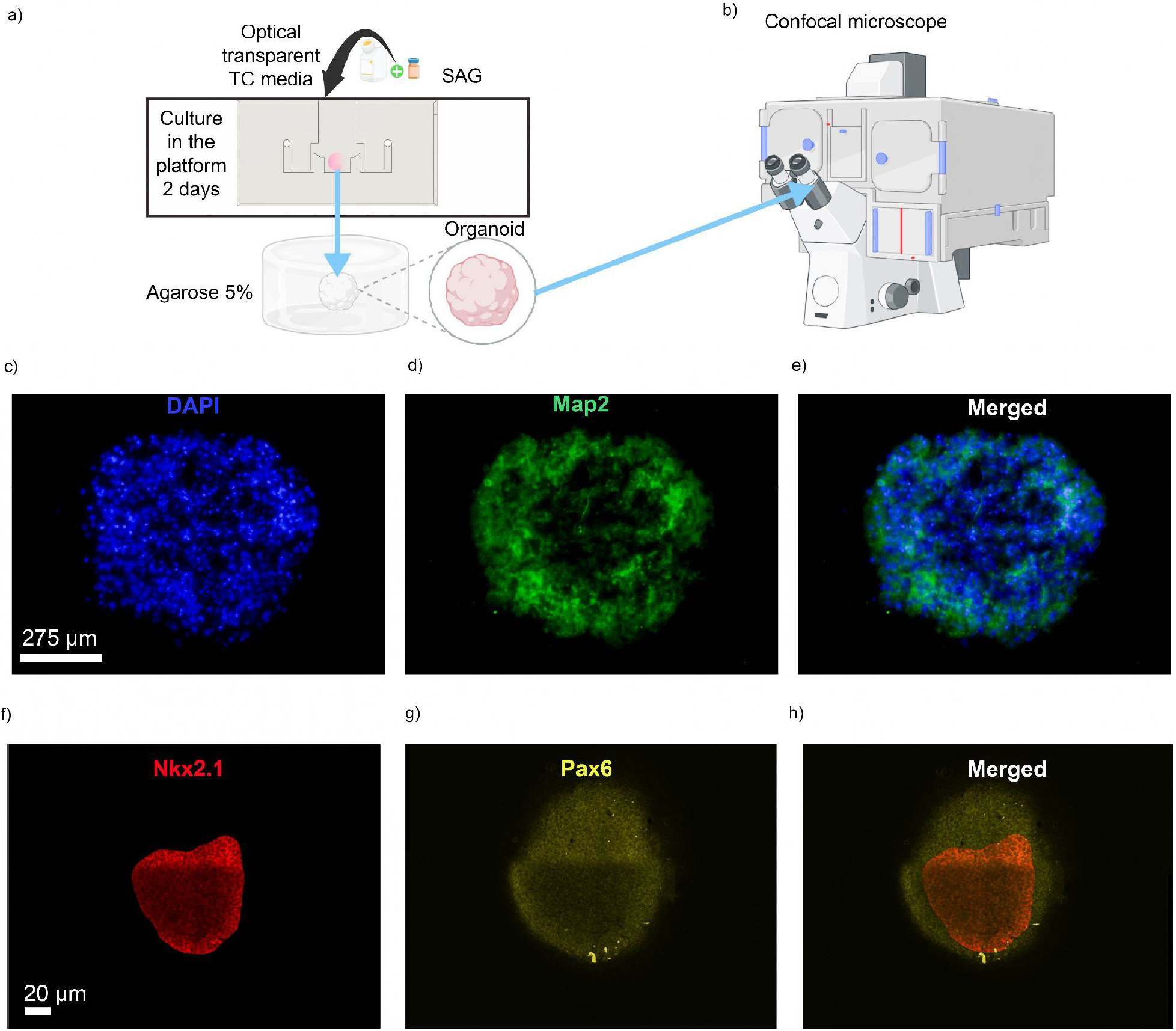
Experimental workflow and immunofluorescent analysis of regionalized cerebral organoids from the platform. (a) Organoids were cultured for 2 days on the microfluidic platform in optically transparent tissue culture (TC) medium supplemented with SAG, then embedded in 5% agarose for imaging. (b) Embedded organoids were imaged using a confocal microscope to acquire volumetric datasets. (c-e) Representative whole-organoid immunofluorescence images showing nuclear staining with DAPI (c, blue), neuronal differentiation marked by Map2 (d, green), and the merged image (e), illustrating global neuronal organization. (f-h) Higher-resolution imaging of regional identity markers showing Nkx2.1 (f, red), Pax6 (g, yellow), and the merged panel (h), revealing spatially segregated ventral (Nkx2.1) and dorsal (Pax6) domains within a single organoid. Scale bars: 275 μm (c-e) and 40 μm (f-h).

Whole-organoid staining showed widespread DAPI nuclear labeling and Map2 expression, indicating global neuronal organization throughout the tissue. Higher-resolution imaging of regional identity markers demonstrated clear spatial segregation between ventral and dorsal domains: Nkx2.1, a marker of ventral forebrain identity, localized separately from Pax6, a marker of dorsal forebrain identity, within the same organoid. The mutually exclusive distribution of these markers is consistent with the establishment of ventral and dorsal patterning and aligns with the CFD simulations, indicating that controlled SAG delivery in the microfluidic system is sufficient to induce organized regional identity across the organoid volume.

To further characterize this patterning, we performed orthogonal reconstruction and spatial analysis of dorsal-ventral organization across three conditions (Fig. 5 and 6). In the first control, organoids cultured with SAG in a standard incubator (Fig. 5a) showed no dorsal-ventral segregation, with Pax6 and Nkx2.1 signals diffusely distributed across orthogonal planes, indicating that chemical induction alone is insufficient for spatial patterning without environmental control. In the second control, organoids cultured without SAG in a standard incubator (Fig. 5n) displayed exclusively dorsal patterning, with Pax6 expression visible across transverse, frontal, and sagittal planes and a complete absence of Nkx2.1 signal, confirming that ventral identity requires SAG treatment. In contrast, organoids maintained in the automated platform with SAG (Fig. 6) developed clear dorsal-ventral segregation, with orthogonal optical sections sampled at multiple depths along the z, x, and y axes confirming consistent marker distribution throughout the entire organoid volume, rather than confined to a surface layer.

**Figure 5.**
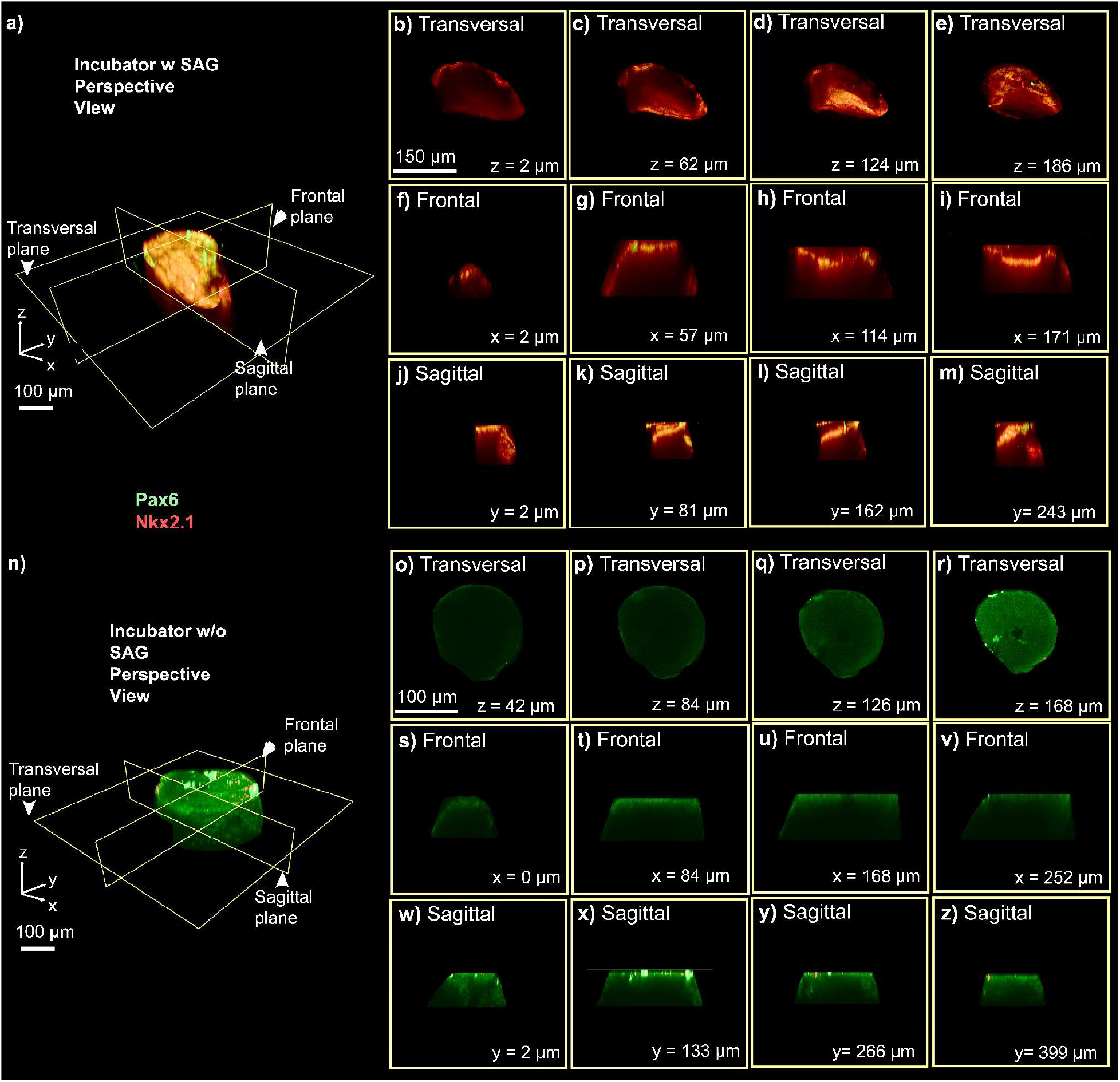
Orthogonal reconstruction and spatial analysis of dorsal-ventral patterning in single organoids cultured under different incubator conditions. (a) Schematic representation of the imaging planes used for orthogonal reconstruction (transversal, frontal, and sagittal), highlighting spatially segregated Pax6 (green, dorsal) and Nkx2.1 (red, ventral) domains within the organoid culture in the incubator with SAG. (b-e) Representative transversal optical sections at increasing z-depths (z = 2 μm, 62 μm, 124 μm, 186 μm) showing the distribution of dorsal and ventral markers along the organoid’s axis in the SAG-treated microfluidic platform. (f-i) Representative frontal optical sections at increasing x-depths (x = 2 μm, 57 μm, 114 μm, 171 μm). (j-m) Representative sagittal optical sections at increasing y-depths (y = 2 μm, 81 μm, 162 μm, 243 μm). (n) Schematic representation of the imaging planes used for orthogonal reconstruction (transversal, frontal, and sagittal), highlighting spatially segregated Pax6 (green, dorsal) and Nkx2.1 (red, ventral) domains within the organoid culture in the incubator without SAG. (o-r) Representative transversal optical sections at increasing z-depths (z = 42 μm, 84 μm, 126 μm, 168 μm) showing the distribution of dorsal and ventral markers along the organoid’s axis in the SAG-treated microfluidic platform. (s-v) Representative frontal optical sections at increasing x-depths (x = 0 μm, 84 μm, 168 μm, 252 μm). (w-z) Representative sagittal optical sections at increasing y-depths (y = 2 μm, 133 μm, 266 μm, 399 μm).

**Figure 6.**
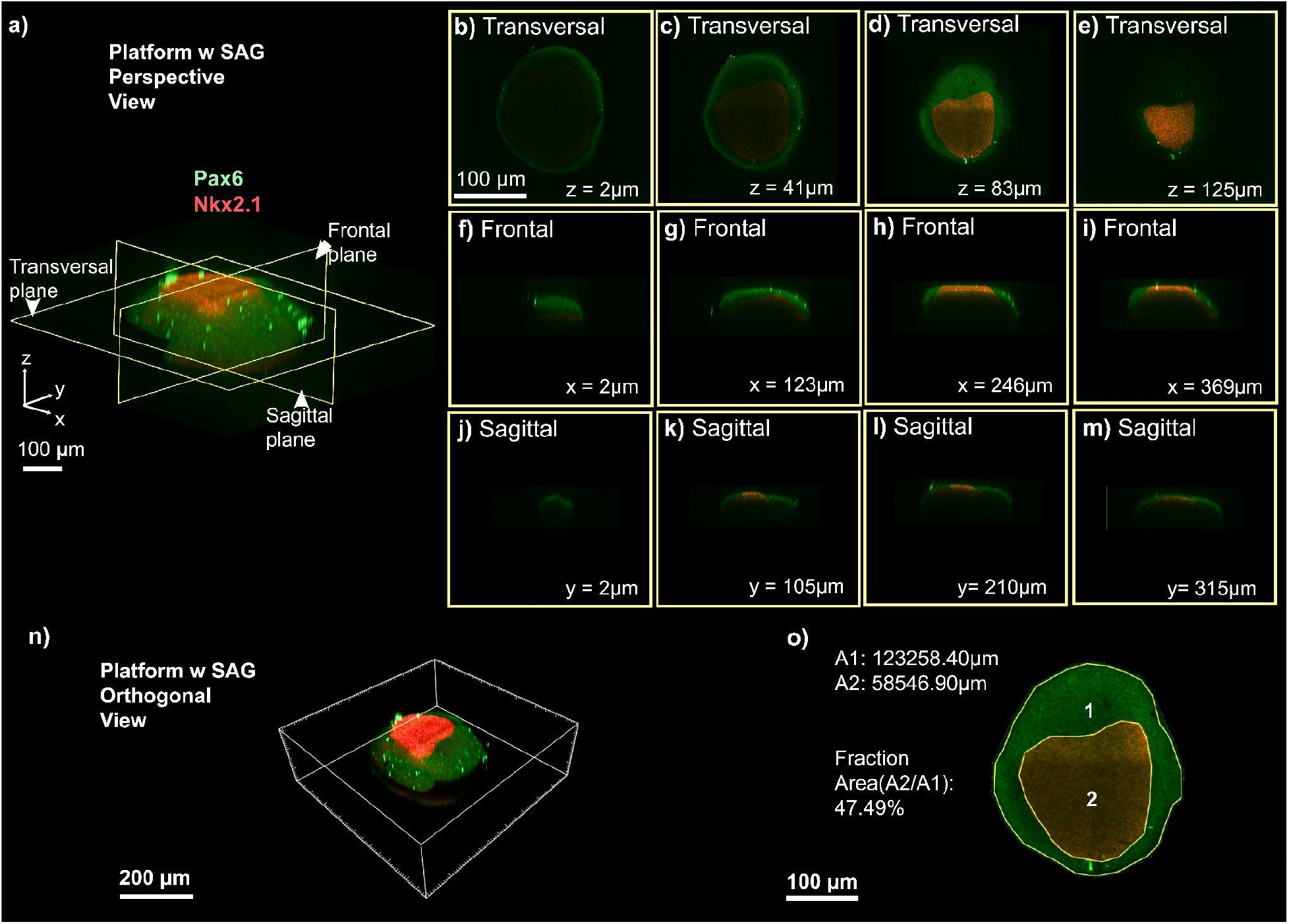
Orthogonal reconstruction and spatial analysis of dorsal-ventral patterning in single organoids cultured under the platform conditions. (a) Schematic representation of the imaging planes used for orthogonal reconstruction (transversal, frontal, and sagittal), highlighting spatially segregated Pax6 (green, dorsal) and Nkx2.1 (red, ventral) domains within the organoid. (b-e) Representative transversal optical sections at increasing z-depths (z = 2 μm, 41 μm, 83 μm, 125 μm) showing the distribution of dorsal and ventral markers along the organoid’s axis in the SAG-treated microfluidic platform. (f-i) Representative frontal optical sections at increasing x-depths (x = 2 μm, 123 μm, 246 μm, 369 μm). (j-m) Representative sagittal optical sections at increasing y-depths (y = 2 μm, 105 μm, 210 μm, 315 μm). (m) Three-dimensional orthogonal rendering of an organoid exposed to SAG within the platform, demonstrating a defined ventral Nkx2.1 region. (n) Quantification of ventral area relative to total organoid area (A1 and A2), illustrating robust ventralization under SAG exposure.

Spatial analysis supported these observations. Fluorescence-based segmentation of organoid cross-sections identified two major regions: an outer domain (A1, 123,258.40 μm^2^) corresponding to the organoid boundary, and a more intense region of ventral marker signal (A2, 58,546.90 μm^2^), representing 47.49% of the total area (Fig. 6o). For visualization purposes, organoids were positioned with the Nkx2.1-positive region centrally to facilitate identification of the ventral domain. A lower-intensity peripheral zone surrounded this region, producing a gradient of marker intensity consistent with partial and localized ventralization. The larger Pax6-positive area is expected, as dorsal identity represents the default fate under basal patterning medium. Apparent overlap at domain boundaries corresponds to a biologically expected transition zone between dorsal and ventral territories, consistent with in vivo forebrain patterning. These quantitative measurements complement the 2D immunostaining analyses (Fig. 4) and confirm that SAG exposure in the automated platform induces robust and spatially organized ventral specification.

## DISCUSSION

Precise control of fluidic patterns and molecular gradients contributes directly to the anatomical shaping of brain organoids through morphogen signaling^33,34,35,36,37,38^. In early brain development, morphogens such as Shh^39^, Wnt^40,41^, or Bmp^42,43^ guide cells toward specific fates by establishing concentration-dependent gradients that define dorsal, ventral, and medial domains^44,45^. Replicating these processes *in vitro* requires not only maintaining organoid viability, but also recreating the spatial and temporal precision of morphogen delivery^46^. The described platform enables localized, tunable delivery of signaling molecules while preserving culture stability, thereby providing an experimental framework for modeling how gradients drive tissue regionalization and linking controlled fluidic environments to observable anatomical outcomes. Validation through simulations and dye-distribution experiments demonstrates that device geometry and organoid presence can influence gradient formation, underscoring the importance of microenvironmental variables in shaping brain architecture.

The platform enables the simultaneous development of multiple brain regions within a single continuous organoid, growing an integrated assembloid while maintaining overall tissue organization and structural integrity. Most existing approaches rely on fusing independently generated organoids, which allows pairing of tissue types but limits spatial coordination, whereas adjacent territories in the embryonic brain emerge, interact, and mature together. By generating two patterned brain regions concurrently, the platform guides region-specific differentiation without requiring later mechanical fusion. Because it orchestrates growth rather than assembly^5,6,45^, the system provides a foundation for scaling to more complex neurodevelopmental interfaces and potentially forming multi-regional brain tissues.

Our experiments used mouse embryonic stem cells (mESCs), which undergo neural induction and fate specification considerably faster than human iPSC-derived organoids. A 2-day SAG exposure (days 3-5) is sufficient to initiate Shh-dependent ventral fate specification in mouse neural progenitors, consistent with prior studies. At this stage, Nkx2.1 and Pax6 expression reflect early progenitor identity rather than fully differentiated cell types. This timing captures the onset of ventral and dorsal fate specification, before interneurons derived from the ventral forebrain begin their rapid migration into dorsal regions during forebrain assembly.

Beyond developmental studies^47,48^, this approach has broader implications for disease modeling and translational applications^49,50,51,52,53^. Many neurodevelopmental disorders^54,55,56,^ such as cortical malformations^57^, arise from disrupted morphogen gradients during early brain formation^2,34,58^. By providing precise control over morphogen delivery, the platform enables researchers to replicate specific pathological conditions and examine their effects on tissue patterning. Combining spatially restricted stimulation^41,59,60^ with longitudinal fluorescence tracking further enables observation of how alterations in gradient dynamics affect cell fate decisions over time, making the platform a versatile tool for investigating the mechanisms underlying neurodevelopmental disorders.

The platform has several limitations. Based on CFD simulations (Fig. 2) and dye experiments (Fig. 3), it establishes an asymmetric SAG gradient across the organoid, with morphogen diffusing inward from the perfused surface over time. We do not have a direct spatial correlation between the predicted gradient and Nkx2.1 distribution, which represents a current limitation. The platform primarily delivers cues to the organoid surface, which may limit penetration in larger tissues. Human organoid samples should be tested to evaluate long-term culture capabilities, and scalability to more complex multi-regional architectures remains untested. Future work should incorporate live morphogen reporters or spatially resolved transcriptomics to map fate specification relative to the delivered gradient more precisely. Despite these limitations, the specificity of fate induction is supported by the mutually exclusive distribution of Nkx2.1 and Pax6 and by the absence of Nkx2.1 signal in SAG-untreated control organoids (Fig. 5). These constraints highlight the need for improvements in gradient complexity, deeper tissue access, and broader biological validation to model human neurodevelopment more faithfully.

## CONCLUSIONS

Our platform enables controlled delivery of molecules to organoid surfaces while maintaining tissue viability, supporting spatially restricted stimulation and longitudinal monitoring. Morphogen gradients can guide the formation of multi-regional organoids, providing a framework for studying neurodevelopmental processes and disorders. The system also allows delivery of macromolecules or antibodies to probe tissue-specific responses and cell-fate decisions. Integration with immunostaining or genetically encoded reporters permits tracking of spatial patterning within organoids. These results show that the platform provides precise control over organoid microenvironments and supports reproducible spatial organization.

## MATERIALS AND METHODS

### Platform fabrication

The platform was designed to support and monitor 3D tissue cultures, and has been previously described in Torres-Montoya et al.(2026)^25^ (Fig. 1). Its primary components include a servo-controlled fluidic pump, a temperature-control unit equipped with an aluminum heating pad, an environmental sensor module, and a microscope with integrated lighting^25^. A fluorescence imaging chamber was also part of the platform, comprising a PDMS-glass microfluidic chip and a microscope. The chip was connected to fluidic lines for media flow using a syringe pump (TECAN Cavro^®^), and an LED ring (NeoPixel, Adafruit) provided illumination. A schematic of the entire setup and its components is shown in Fig. 1d. Complementary views of the platform are shown in supplementary Figs. 4 and 5.

The PDMS-glass microfluidic chip was fabricated using a single microscope slide and 6 grams of silicone. See supplementary Fig. 6 for the manufacturing details. The fabrication process began with printing a solid mold from Model V2 resin on a Form 3 SLA printer (Formlabs). The printed mold was cleaned in isopropanol (IPA) using an ultrasonic bath for 20 minutes, dried under nitrogen, and then cured under UV light (405 nm) at 60°C for 40 minutes. Then, PDMS (Sylgard 184, Dow Corning) was mixed with the curing agent in a 10:1 weight ratio. This mixture was degassed in a vacuum chamber for 2 hours and then cured in an oven at 60°C for 6 hours. A surface activation step was performed to bond the PDMS to a 25 mm × 75 mm microscope glass slide (Fisherbrand™ Superfrost™, Fisher) using O2 plasma reactive ion etching (RIE). The PDMS and glass were then aligned and pressed together to form a tight seal^25^. The assembled bio-chamber was placed in a vacuum oven for an additional six hours at 60°C, with 10 grams of pressure applied to enhance adhesion between the layers.

### Imaging Acquisition

Imaging was performed using the Dino-Lite Edge AM4115T-GRFBY digital microscope, which has a resolution of 1.3 megapixels and a frame rate of 30 frames per second (FPS). The microscope is equipped with two LED light sources for fluorescent imaging: one set of four blue LEDs (excitation at 480 nm) and another set of four yellow LEDs (excitation at 575 nm). The LED sets could be easily switched using the built-in software, providing flexibility in imaging. The corresponding emission wavelengths were 510 nm for the blue LEDs and 610 nm for the yellow LEDs. The microscope provides magnification up to 220x, enabling close-up visualization of samples. Video recordings were made using Videocapture 2.0 (DinoEdge, Dino-Lite), a free software provided by the manufacturer^61^. This software supported smooth video capture and efficient data management during experiments.

### Computational Fluid Dynamics

To evaluate the platform design, computational fluid dynamics (CFD) simulations were conducted using COMSOL Multiphysics 5.5 (Stockholm, Sweden), and a previously reported protocol was employed^25^. The goal was to simulate fluid behavior under controlled conditions. The model assumed a linear flow speed of 5×10^−3^ms^−1^ and a 70 μL media injection into a well with a depth of 5 mm and a diameter of 5.6 mm. Water properties were used to represent the liquid phase, while air properties were used to model the gas phase. The simulations were conducted under incubator conditions, with water density set to 997 kg/m^3^ and viscosity to 6.92 × 10^−3^ kg/(m·s). The simulation included the spread of the CellTracker Green BODIPY Dye (Thermo Fisher Scientific, #C2102), a commonly used live-cell membrane stain. The dye was introduced at a concentration of 50 nmol/m^3^ at the inlet as a boundary condition, and its diffusion across the organoid surface was modeled using a diffusion coefficient of 6.5×10^-14^ m^2^/s, as previously describedt^62^. The platform was maintained at a reference temperature of 37°C. Fluid flow was governed by the Navier-Stokes equations, with boundary conditions established from earlier experiments conducted in shaker environments. To accurately capture the interface between the air and liquid phases, the level set method was applied. This approach enabled a smooth transition at the interface, with the level-set function ranging from 0 (air) to 1 (liquid); a midpoint value of 0.5 was chosen to represent a balanced surface-tension condition and to ensure computational efficiency. The model geometry was bounded by solid walls, with a non-slip condition applied to all surfaces except the top, which was open to incubator air and set with a slip condition. The simulation mesh consisted of 519,830 tetrahedral elements. A spherical organoid shape (1.8 mm diameter) was introduced, based on imaging data from day 13 of culture. Environmental conditions were set to mimic those of an incubator: 1 atm, 37°C, and a gas mixture of 5% CO2, 17% O2, and 78% N2. To confirm the simulation’s reliability, a mesh independence test was performed. Final results, including flow-velocity fields, were visualized as streamlines overlaid on a central cross-sectional plane.

### Experimental Verification of the CFD

We used the CellTracker Green BODIPY Dye (Thermo Fisher Scientific #C2102) in conjunction with a PDMS/Glass biocompatible chip to examine the presence and dynamics of velocity fields in cortical organoids. On day five of the experiment, we used a programmable pump to set a constant velocity of 5×10^−3^ms^−1^. The inlet and outlet volumes were set to 53 μL and 50 μL, respectively, with a recirculation time of 150 seconds. For real-time imaging, we used the Dino-Lite Edge AM4115T-GRFBY, a digital microscope with a 1.3 MP resolution and a frame rate of 30 frames per second (fps). The microscope’s green LEDs were used for fluorescent imaging of the objects. This experimental setup was replicated across three organoids, enabling comprehensive observations and analysis. Subsequently, a fluorescence-mediated pump test was conducted to verify the functionality and reliability of the microfluidic chamber experimentally. This test involved pumping a fluorescence-labeled medium through the platform and observing its behavior under controlled conditions. The objective was to ensure that fluid flow remained uniform, without leakage or undesirable disturbances, while maintaining the desired characteristics of the sample medium. The image intensities were analyzed using the ImageJ pixel processing module, and LED microscope intensity was measured with a digital lux meter (LX1332B, Dr. Meter). These measures were correlated to obtain intensity image values, which were then converted to concentration values using a calibration curve (see Supplementary Figures 7 and 8).

### Embryonic Stem Cell Line Maintenance

All experiments utilized the BRUCE4 mouse embryonic stem cell (ESC) line (C57BL/6 background). Before experimentation, the ESCs were tested for mycoplasma contamination. ESCs were cultured on plates coated with Recombinant Protein Vitronectin (Thermo Fisher Scientific # A14700). The maintenance medium for mESCs consisted of Glasgow Minimum Essential Medium (Thermo Fisher Scientific # 11710035), supplemented with Embryonic Stem Cell-Qualified Fetal Bovine Serum (Thermo Fisher Scientific # 10439001), 0.1 mM MEM Non-Essential Amino Acids (Thermo Fisher Scientific # 11140050), 1 mM Sodium Pyruvate (Millipore Sigma # S8636), 2 mM Glutamax supplement (Thermo Fisher Scientific # 35050061), 0.1 mM 2-Mercaptoethanol (Millipore Sigma # M3148), and 0.05 mgmL^-1^ Primocin (Invitrogen # ant-pm-05). The mESC maintenance media were supplemented with 1,000 units mL^-1^ of Recombinant Mouse Leukemia Inhibitory Factor (Millipore Sigma # ESG1107). The culture medium was changed daily to maintain optimal conditions for ESC growth. ReLeSR passaging reagent (Stem Cell Technologies # 05872) was used according to the manufacturer’s instructions for cell dissociation and passaging. To preserve cells, ESCs were frozen in mFreSR cryopreservation medium (Stem Cell Technologies #05855) according to the manufacturer’s guidelines.

### Generation of Cortical Organoids

Mouse embryonic stem cells (mESCs) were first dissociated into single cells using TrypLE Express (Thermo Fisher #12604021) for 5 minutes at 37°C, following previously published protocols ^31^. The cells were then re-aggregated in Lipidure-coated 96-well V-bottom plates at a density of 3,000 cells per well, suspended in 150 μL of mESC maintenance medium. This medium was supplemented with 10 μM ROCK inhibitor Y-27632 (Tocris #1254) and 1,000 U/mL recombinant mouse LIF (Millipore Sigma #ESG1107) to support cell survival and self-renewal. After 24 hours, the medium was replaced with a dorsal forebrain patterning medium. This medium consisted of DMEM/F12 with GlutaMAX (Thermo Fisher #10565018), 0.1 mM MEM Non-Essential Amino Acids (Thermo Fisher #11140050), 1 mM sodium pyruvate (Sigma #S8636), 1X B-27 minus Vitamin A (Thermo Fisher #12587010), 1X Chemically Defined Lipid Concentrate (Thermo Fisher Scientific # 11905031), and 0.05 mg/mL Primocin (InvivoGen #ant-pm-05). To guide the cells toward a dorsal forebrain fate, the medium was further enriched with 10 μM Y-27632, 5 μM WNT inhibitor XAV939 (StemCell Technologies #100-1052), and 5 μM TGF-β inhibitor SB431542 (Tocris #1614). Media were changed daily, and both B-27 and CD lipid concentrate supplements were added after filtration to preserve their hydrophobic components.

On day 5, the aggregates were transferred to ultra-low-attachment plates (Sigma #CLS3471) containing fresh neuronal differentiation medium and placed on an orbital shaker at 68 rpm to enhance mixing and prevent clumping. From day 6 to day 12, the progenitor expansion medium included DMEM/F12 with GlutaMAX, BrainPhys Neuronal Medium (StemCell Technologies #05790), 1X B-27 minus Vitamin A, 1X CD Lipid Concentrate, 0.1 mM MEM NEAA, 0.05 mg/mL Primocin, and 200 μM ascorbic acid (Sigma #49752). Cultures were kept in a 5% CO2 incubator, and the medium was replaced every other day. Starting on day 15, the organoids were transitioned to a neural maturation medium. This included BrainPhys Neuronal Medium supplemented with 1X B-27 Plus (Thermo Fisher #A3582801), 1X CD Lipid Concentrate, 5 μg/mL heparin (Sigma #H3149), and 0.05 mg/mL Primocin. Ascorbic acid (200 μM) was included in the medium until day 25. Media changes continued every 2 to 3 days. To prevent organoid fusion, cultures (16 organoids per well) were maintained on a shaker at 60 rpm.

### Cell migration and organoid regionalization

For localized patterning and migration experiments, mouse embryonic stem cells (mESCs) were re-aggregated in Lipidure-coated 96-well V-bottom plates at a density of 25,000 cells per well, for imaging resolution purposes, in 150 μL of maintenance medium supplemented with 10 μM ROCK inhibitor (Y-27632) and 1,000 U/mL recombinant mouse LIF. After 24 hours, the medium was replaced with dorsal forebrain patterning medium containing 10 μM Y-27632, 5 μM WNT inhibitor (XAV939), and 5 μM TGF-β inhibitor (SB431542), and the medium was changed daily. On day 3, a subset of organoids was transferred into the microfluidic platform designed for localized delivery of signaling molecules. The injected medium contained the same components as the dorsal patterning medium, supplemented with 100 nM Sonic Hedgehog agonist (SAG; Millipore Sigma #SIAL-SML1314), enabling regionalized ventralization within the targeted area while the remaining portion of the organoid continued under dorsal patterning conditions. Control groups included organoids cultured under uniform dorsal conditions and those exposed to 100 nM SAG throughout the entire medium. Media were refreshed daily for all conditions. On day 5, organoids were fixed for immunostaining and imaging analyses to assess spatial distribution, migration behavior, and regional marker expression.

### Immunochemistry and immunostaining

For immunofluorescence staining, organoids were collected, fixed in 4% paraformaldehyde (Thermo Fisher Scientific #28908), and cryoprotected in 30% sucrose solution. Samples were embedded in a 1:1 mixture of Tissue-Tek O.C.T. Compound (Sakura #4583) and 30% sucrose solution and sectioned at 12 μm using a cryostat (Leica Biosystems #CM3050). Sections were washed twice for 5 minutes each in 1× PBS, followed by a single wash in deionized water (Chem World #CW-DW-2G). Sections were then incubated for 1 hour in blocking solution containing 5% (v/v) donkey serum (Millipore Sigma #D9663) and 0.1% Triton X-100 (Millipore Sigma #X100). Primary antibodies were applied and incubated overnight at 4°C. The sections were washed three times for 10 minutes in PBS and incubated with secondary antibodies for 90 minutes at room temperature. After three additional 10-minute washes in PBS, sections were mounted using Fluoromount-G Mounting Medium (Thermo Fisher Scientific #00-4958-02). Primary antibodies included mouse and rabbit anti-Map2 (Proteintech #17490-1-AP) and were used at 1:2,000. Alexa-conjugated secondary antibodies (Thermo Fisher Scientific) were used at a 1:750 dilution. Nuclear counterstaining was performed with 300 nM DAPI (4′,6-diamidino-2-phenylindole, dihydrochloride) (Thermo Fisher Scientific #D1306). Imaging was performed using EVOS-FL Auto-2 fluorescence microscope (Thermo Fisher Scientific) equipped with a TRITC filter set (Ex/Em = 540/590 nm).

For whole-organoid immunofluorescence staining, mouse brain organoids were fixed at room temperature for 45 minutes in 4% paraformaldehyde. After fixation, they were washed 3 times in PBS and stored at 4°C. For whole-organoid immunostaining, the organoids were blocked for 24 hours at room temperature in PBS supplemented with 0.2% gelatin (VWR, 24350.262) and 0.5% Triton X-100 (Millipore Sigma, X100) (PBSGT). Samples were then incubated with rabbit anti-Nxk2.1 (Abcam, ab76013, 1:100) and mouse anti-Pax6 (BD Biosciences, 561462, 1:100) primary antibodies for 7 days at 37 oC at 70 rpm in PBSGT + 1 mg/ml Saponin Quillaja sp (Sigma Aldrich, S4521) (PBSGTS). Following primary antibody incubation, samples were washed 6 times in PBSGT at room temperature for 1 day. Secondary antibody staining was done using the Alexa Fluor 488 Donkey anti-mouse (1:250) and the Alexa Fluor 546 Donkey anti-rabbit (1:250) antibodies, and the nucleus was stained using DAPI dye. Secondary and nuclear staining were performed for 1 day at 37 °C at 70 rpm in PBSGTS. Samples were then washed 6 times in PBSGT at room temperature for 1 day.

### Confocal imaging

Organoid imaging was performed using a Zeiss 880 confocal microscope equipped with Airyscan Fast, and 4x magnification is shown in Fig.4 (a-c) and Supplementary Fig. 9. Image acquisition was performed using Zen Blue software, and Z-stacks (n=100) were acquired with a 1.53 μm spacing from three non-adjacent cryosections per organoid. Tile scanning was applied when the organoid exceeded a single field of view. Two organoid replicates were analyzed per cell line and condition (dorsal/ventral) across two independent cell lines (ES-E14TG2a and KH2). Raw image files (.czi) were converted to.ims format using Imaris (version 10.2) and then deconvolved with AutoQuant X3 3.1 before further analysis in Imaris (version 10.2). Nucleus segmentation was first performed on the DAPI channel, followed by spot detection with an XY diameter of 4.5 μm, based on average cell diameters. For marker quantification, Pax6 and Nkx2.1 cells were identified using the same spot-detection parameters, with additional colocalization constraints requiring a maximum distance of 14 μm from the DAPI signal centers. The workflow was automated through Imaris Arena to ensure parameter consistency across all patterning conditions. Quantitative outputs included absolute counts of total DAPI nuclei, Pax6/DAPI double-positive cells, and Nkx2.1/DAPI double-positive cells.

### Platform Sterilization

To ensure a sterile environment and prevent contamination, all platform components were subjected to rigorous sterilization before each experiment. The PDMS/glass chip, temperature feedback sensor, thermocouple, media tubing, and programmable pump were immersed in a high-level disinfectant (CIDEX OPA, ASP) for 20 minutes. After disinfection, these components were thoroughly rinsed with ultrapure distilled water (Invitrogen Ultra-Pure, ThermoFisher) for an additional 20 minutes to remove any residual disinfectant. Surfaces and accessories—including the chamber, chip holder, environmental sensor, power cable, and microscope—were wiped with hydrogen peroxide solution (Oxivir TB, Diversey) for 1 minute. These stringent procedures ensured comprehensive decontamination and maintained sterile conditions throughout the study.

## DATA AVAILABILITY

All custom scripts, feeding rates, 3D printed files, microscope images, and CFD videos have been made available at [https://github.com/sebtomon89/braingeneersdifussionproject]. Additional modified scripts are available upon request. All other relevant data are available from the corresponding author on request.

## CODE AVAILABILITY

Details of publicly available software used in the study are given in the “Data availability” section. Apart from this, no unique custom code or mathematical algorithms were central to reaching the conclusions of this work.

## ACKNOWLEDGMENTS

This work was supported by Schmidt Futures (SF857 to S.R.S., D.H., and M.T.); the National Human Genome Research Institute (RM1HG011543 to S.R.S., D.H., and M.T.); the National Science Foundation (NSF; 2134955 to S.R.S., D.H., and M.T.; 2034037 to M.T.; 2515389 to D.H., M.A.M.-R., and M.T.); the National Institute of Mental Health (U24MH132628 to D.H. and M.A.M.-R.); the National Institute of Neurological Disorders and Stroke (U24NS146314 to D.H. and M.A.M.-R.); the California Institute for Regenerative Medicine (DISC4-16285 to S.R.S., M.A.M.-R., and M.T.; DISC4-16337 to M.A.M.-R.); the University of California Office of the President (M25PR9045 to S.R.S., M.A.M.-R., and M.T.); and the Brain and Behavior Research Foundation (33184 to M.A.M.-R.). S.V.-C. was partially supported by the Graduate Pedagogy Fellowship from the University of California Santa Cruz Teaching and Learning Center.

In addition, we thank the UCSC Life Sciences Microscopy Center, RRID: SCR_021135, for providing the confocal microscope to acquire the images. During the preparation of this work, the authors used ChatGPT and Grammarly to improve clarity and sentence structure. After using this tool/service, the authors reviewed and edited the content as needed and assumed full responsibility for the publication. We are thankful to Sebastian Hernandez for his feedback on the organoid preparation protocol for this manuscript.

## AUTHOR CONTRIBUTIONS STATEMENT

S.T.-M., M.A.M.-R., and M.T. conceived the experiments. S.T.-M., S.T.S., and S.V.-C. performed all the experiments. S.R.S., D.H., M.A.M.-R., and M.T. supervised the work and secured funding. S.T.-M., M.A.M.-R. and M.T. wrote the manuscript with contributions from all authors.

