## Supplementary Materials for "Microfluidic Control of Dorsal-Ventral Patterning Within a Single Forebrain Organoid"

### Supplementary Information

**Formulation:**

Computational fluid dynamics formulation.

**Supplementary Tables:**

Supplementary Table 1. CFD Initial Conditions.
Supplementary Table 2: Calibration curve of 50 nM at 0.79 $W/m^2$

**Supplementary Figures**:

Supplementary Figure 1. Computational domain definition, governing equations, boundary conditions, and mesh independence validation for the simulation model.
Supplementary Figure 2. Characterization of dye distribution in microfluidic devices with different geometries.
Supplementary Figure 3. 3D reconstruction of patterned brain organoids cultured within the PDMS platform.
Supplementary Figure 4. Complementary views of the platform.
Supplementary Figure 5. Schematic overview of the automated microfluidic culture platform.
Supplementary Figure 6. Fabrication and testing of the microfluidic device.
Supplementary Figure 7. Quantification of BODIPY dye diffusion and concentration dynamics within the microfluidic chip.
Supplementary Figure 8. Calibration and spatial distribution of pixel intensity across the organoid cross-section.
Supplementary Figure 9. Workflow for organoid immunostaining and high-resolution imaging.

#### Formulation

Three distinct domains were used to simulate the two-phase flow: air, the medium, and the interface between them. The liquid-air interface was treated as a free surface, and the Navier-Stokes equations were employed to model the unsteady three-dimensional mass and momentum within the homogeneous domain, which can be defined as follows:

$$\frac{\partial}{\partial t}(\rho\bar{v}) + \nabla \cdot (\rho\overline{vv}) =- \nabla p + \nabla(\mu\nabla\bar{v}) + \rho\bar{F} \qquad (1)$$

$$\frac{\partial}{\partial t} + \nabla \cdot (\rho\overline{vv}) = 0 \qquad (2)$$

The equations (1) and (2), where ***v*** represents the velocity, ρ denote the density, ***p*** signifies the pressure, and ***F*** represents the external forces acting within each single-phase domain. This equation describes the conservation of momentum. The level-set method was implemented to accurately capture the free-surface flow interface. This method enables the calculation of the advected velocity in the solution of the Navier-Stokes equation. The level-set function Φ is introduced to describe the relationship between the free liquid surface and surface tension. The formulation of the level set method can be expressed as:

$$\frac{\partial}{\partial t} + \nabla\phi \cdot \mu = F \qquad (3)$$

In Equation (3), Φ represents the level set function, ***u*** denotes the velocity, and ***F*** represents the source term that accounts for the influence of surface tension. By using the level-set method, the dynamics of the free surface can be accurately incorporated into the solution of the Navier-Stokes equations.

In this work, we address the numerical solution of a physical problem involving the transport and absorption of a substance, specifically fluorescence dyes. To model this phenomenon, we employ the mass-balance equation, which describes the transport of the diluted dye through the surrounding medium via advection and diffusion. The mass balance equation is given by:

$$\frac{\partial c_i}{\partial t} + \nabla \cdot J_i + u \cdot \nabla c_i = R_i \qquad (4)$$

In Equation 4, $\boldsymbol{C_i}$ represents the concentration of the species in $molm^{-3}$, Di represents the diffusion coefficient in $m^2s^{-1}$, $\boldsymbol{R_i}$ represents the reaction rate in $molm^{-3}s^{-1}$, and ***u*** represents the mass-averaged velocity in $ms^{-1}$. The diffusion coefficient, $\mathbf{D_i}$, is related to the flux density, $\mathbf{J_i}$, in $molm^{-2}s^{-1}$, and is expressed as:

$$J_i =- D\nabla_c \qquad (5)$$

Considering the non-compressibility of the fluid, the advective term in the mass balance equation is formulated in a non-conservative manner, which can be written as:

$$\frac{\partial c}{\partial t} + u \cdot \nabla c = \nabla\cdot c_i + R \qquad (6)$$

**Supplementary Table 1. CFD Initial Conditions.**

| Variable | Value | Units |
|---|---|---|
| **Inlet-Outlet Initial Velocities** | 0.005 | m/s |
| **Diffusion Coefficient** | $6.5x10^{-14}$ | $m^2/s$ |
| **Initial Concentration** | 0.00005 | $mol/m^3$ |
| **Density** | 1000 | $kg/m^3$ |
| **Dynamic Viscosity** | 0.002-0.003 | Pa.s |
| **Porosity** | 0.15 | a.u. |
| **Number of elements** | 519.830 | Elements |

**Supplementary Table 2: Calibration curve of 50 nM at 0.79 $W/m^2$**

| Con (nM) | M1 | | M2 | | M3 | | Std Dev Power | Power Int Avg | Std Dev Pixel | Pixel Int Avg |
|---|---|---|---|---|---|---|---|---|---|---|
| | Pixel Intensity | Power Intensity | Pixel Intensity | Power Intensity | Pixel Intensity | Power Intensity | | | | |
| 0 | 15.577 | 0.641 | 15.783 | 0.649 | 14.906 | 0.613 | 0.019 | 0.634 | 0.458 | 15.422 |
| 5 | 15.940 | 0.656 | 16.194 | 0.666 | 16.090 | 0.662 | 0.005 | 0.661 | 0.128 | 16.075 |
| 10 | 16.303 | 0.671 | 16.129 | 0.663 | 16.229 | 0.668 | 0.004 | 0.667 | 0.087 | 16.220 |
| 15 | 16.666 | 0.686 | 16.438 | 0.676 | 15.677 | 0.645 | 0.021 | 0.669 | 0.518 | 16.260 |
| 20 | 17.029 | 0.700 | 17.323 | 0.713 | 17.809 | 0.733 | 0.016 | 0.715 | 0.394 | 17.387 |
| 25 | 17.391 | 0.715 | 17.926 | 0.737 | 16.954 | 0.697 | 0.020 | 0.717 | 0.487 | 17.424 |
| 30 | 17.754 | 0.730 | 17.143 | 0.705 | 16.678 | 0.686 | 0.022 | 0.707 | 0.540 | 17.192 |
| 35 | 18.117 | 0.745 | 18.078 | 0.744 | 17.223 | 0.708 | 0.021 | 0.732 | 0.505 | 17.806 |
| 40 | 18.480 | 0.760 | 19.250 | 0.792 | 18.725 | 0.770 | 0.016 | 0.774 | 0.393 | 18.818 |
| 45 | 18.843 | 0.775 | 17.978 | 0.740 | 17.935 | 0.738 | 0.021 | 0.751 | 0.512 | 18.252 |
| 50 | 19.206 | 0.790 | 19.235 | 0.791 | 19.381 | 0.797 | 0.004 | 0.793 | 0.094 | 19.274 |

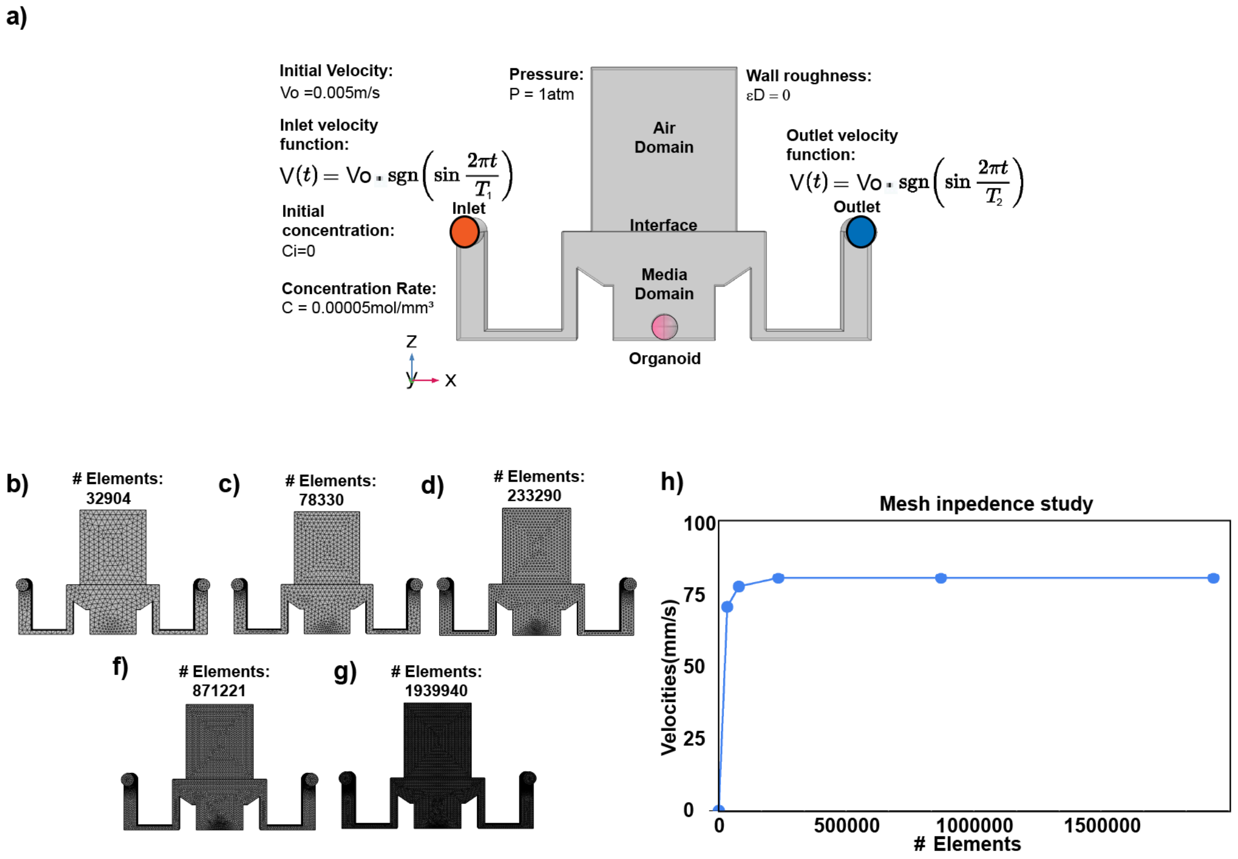


**Supplementary Figure 1. Computational domain definition, governing equations, boundary conditions, and mesh independence validation for the simulation model.** (a) Schematic of the computational domain, comprising two phases: the air domain (top) and the media domain (bottom), separated by an interface. Solute transport occurs within the media domain, governed by volume—of—fluid (VOF)-based Navier-Stokes and interface-tracking equations for two-phase flow. The left side depicts the incompressible momentum conservation and continuity equations, while the right side presents the phase-field equation governing the evolution of the interface. The boundary conditions applied to the inlets, described by time-dependent sign functions, induce anti-phase pulsatile flow on the left and right sides with periods $T_1$ and $T_2$, respectively. (b-g) Meshes used for the grid sensitivity analysis, ranging from coarse to fine, capture geometric features of both the media and air domains. Each mesh is tailored to resolve the interface, inlet curvature, and chamber core. (h) Mesh convergence study. The graph shows that the simulation results stabilize beyond ~400,000 elements, confirming the numerical solution's mesh independence. These results validate the suitability of the mesh resolution used in the main simulations.

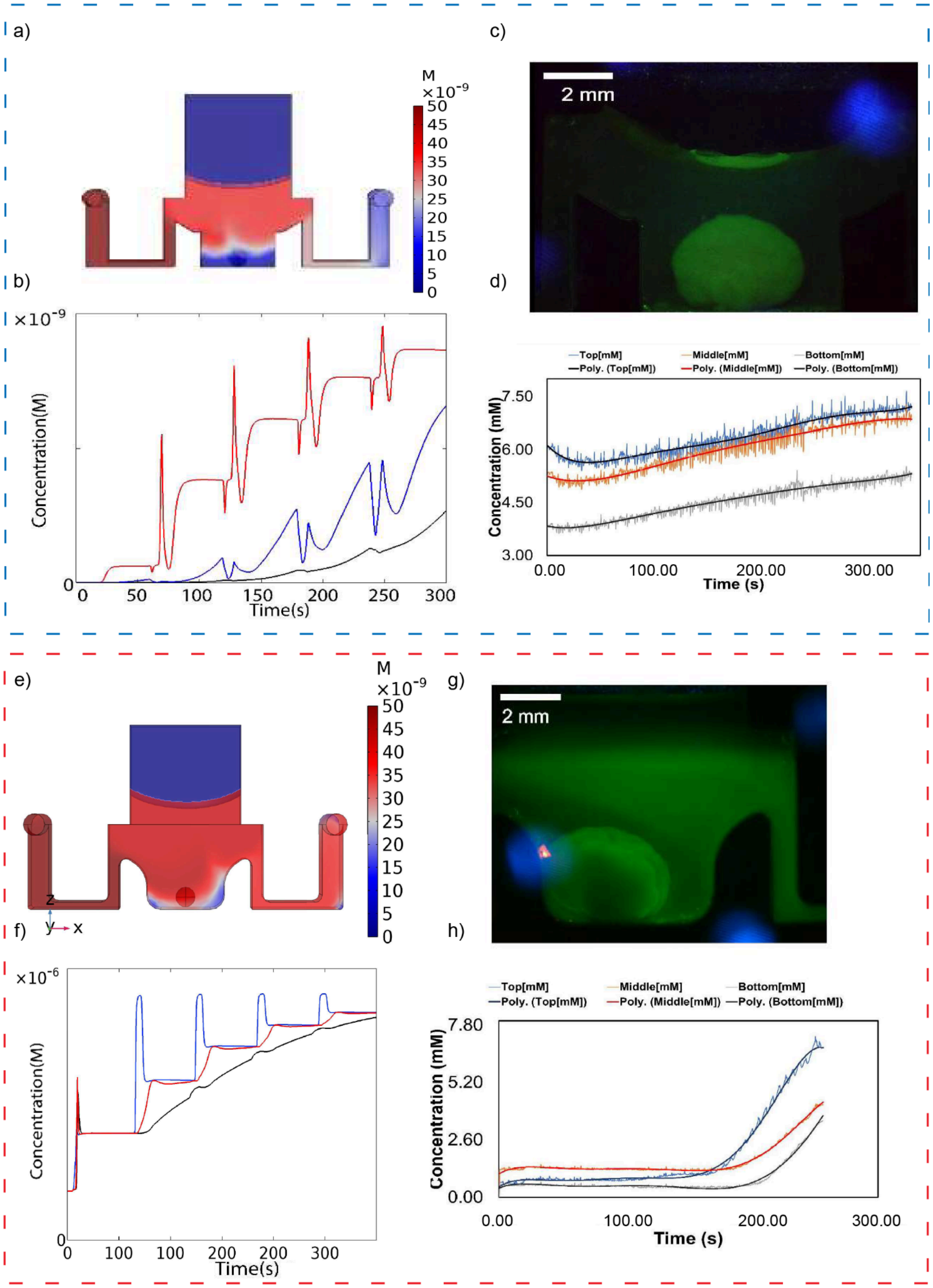


**Supplementary Figure 2. Characterization of dye distribution in microfluidic devices with different geometries**. (a, e) COMSOL simulations of 10 mM BODIPY dye showing concentration fields within a rectangular-bottom chip (a) and a curved-bottom chip (e). (b, f) Simulated temporal concentration profiles across the top, middle, and bottom regions of interest (ROI 1-3), demonstrating geometry-dependent gradient evolution. (c, g) Fluorescence images of the corresponding PDMS chips with BODIPY dye (scale bars: 2 mm). (d, h) Experimental measurements of dye concentration dynamics across ROI 1-3, confirming that geometric shape alters local concentration profiles, consistent with simulation results.

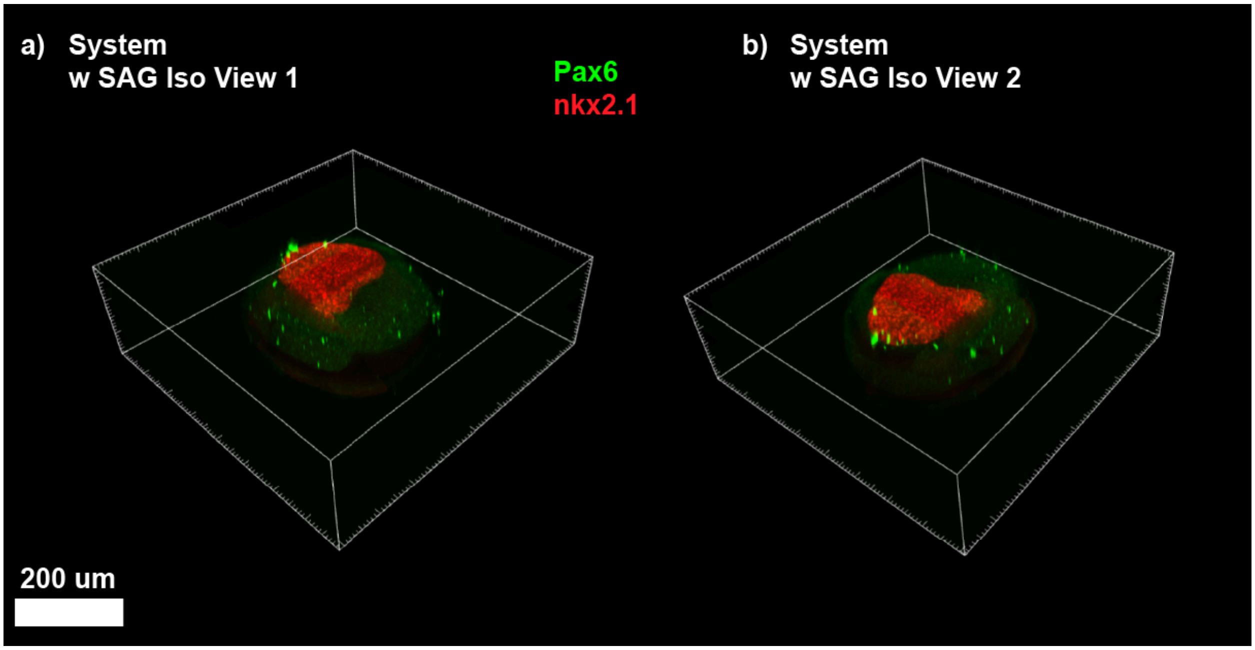


**Supplementary Figure 3. 3D reconstruction of patterned brain organoids cultured within the PDMS platform.** (a-b) Confocal isometric projections showing dorsal (Pax6, green) and ventral (Nkx2.1, red) forebrain marker expression in organoids patterned with SAG. Both views depict the same sample rotated to illustrate marker distribution across the organoid volume. Pax6 expression localized to the periphery, while Nkx2.1 expression was concentrated in a distinct ventral-like region, confirming spatial segregation of forebrain domains within the platform.

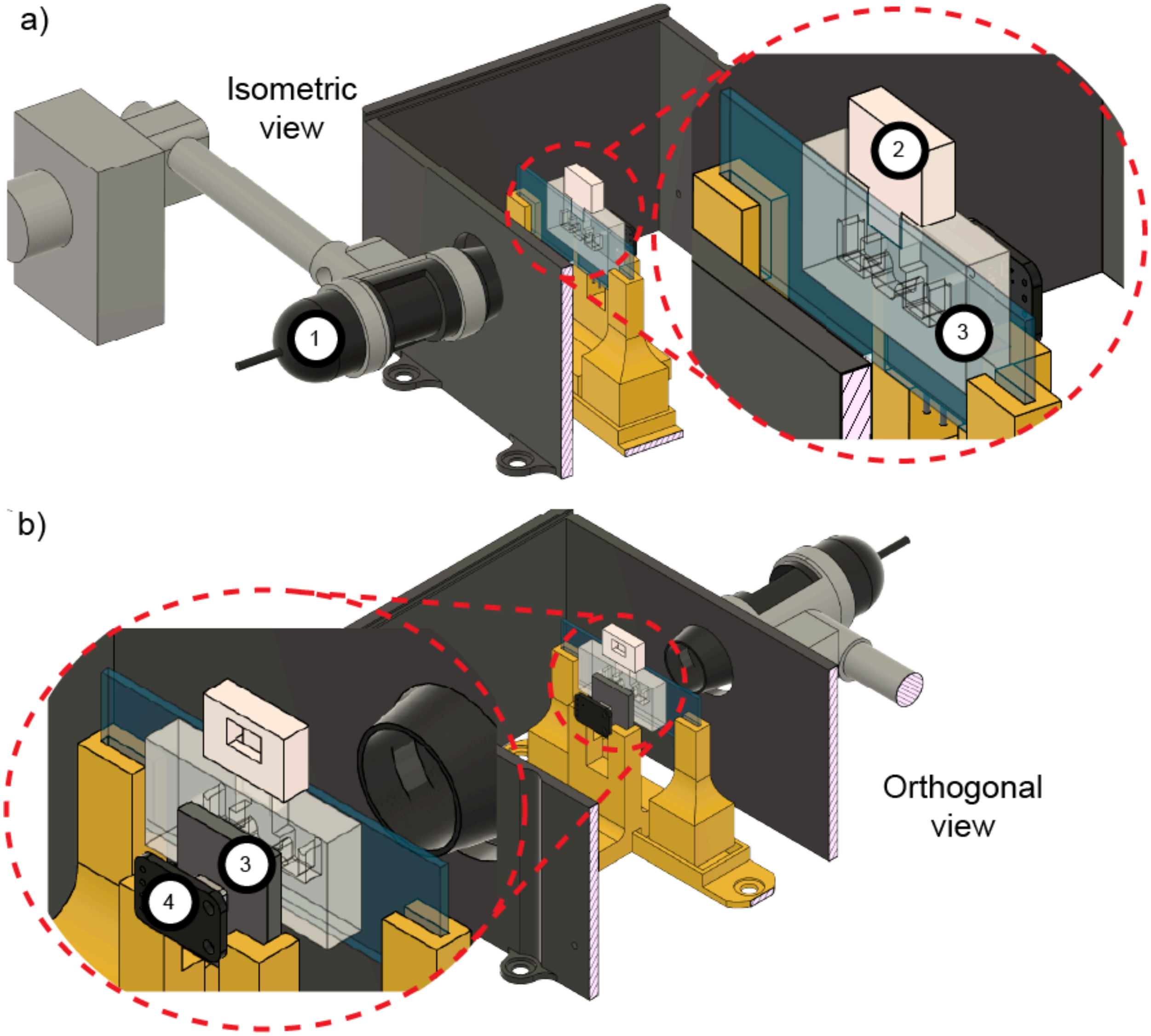


**Supplementary Figure 4. Complementary views of the platform.** Auxiliary isometric views for the platform, a) isometric view, showing the microscope (1), the chip cap (2), and the PDMS/Glass chip (3), and b) where the heat pad(3) and the temperature check sensor (4) are shown.

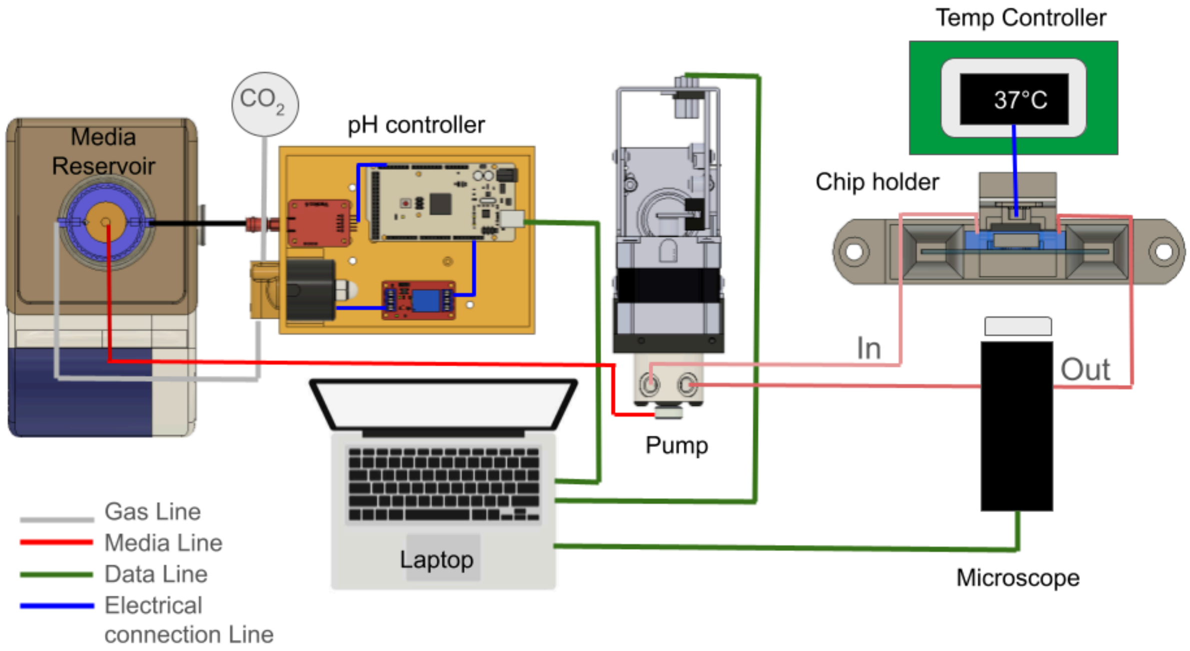


**Supplementary Figure 5. Schematic overview of the automated microfluidic culture platform.** The platform integrates controlled media delivery, environmental regulation, and real-time imaging for microphysiological chip experiments. A media reservoir supplies fresh medium through a gas-regulated ($CO_2$) headspace and feeds the circuit via the pump (red lines, media flow). The pH controller monitors and adjusts the media pH using embedded sensors and electronic control (electrical and data connections are shown in blue and green). The conditioned medium is routed to the chip holder, maintained at 37 °C by an external temperature controller. Effluent exits the chip through the outlet port for downstream analysis. The platform interfaces with a laptop for platform control and data acquisition, while the microscope provides live imaging of the cultured sample. Color-coded lines indicate gas (gray), media (red), data (green), and electrical (blue) connections.

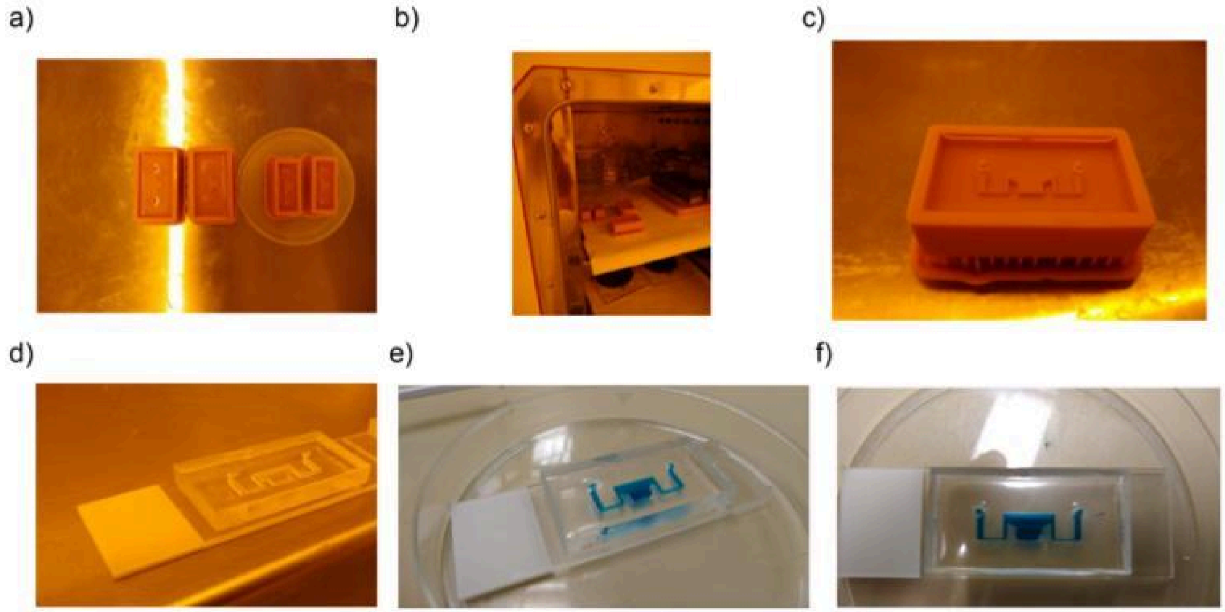


**Supplementary Figure 6. Fabrication and testing of the microfluidic device.** (a) 3D-printed molds prepared for casting**.** (b) Molds are placed inside the incubator for curing. (c) Example of a cured mold ready for device fabrication. (d) PDMS cast after demolding. (e-f) Final microfluidic device bonded to a glass slide, tested with dye to confirm proper channel formation and sealing.

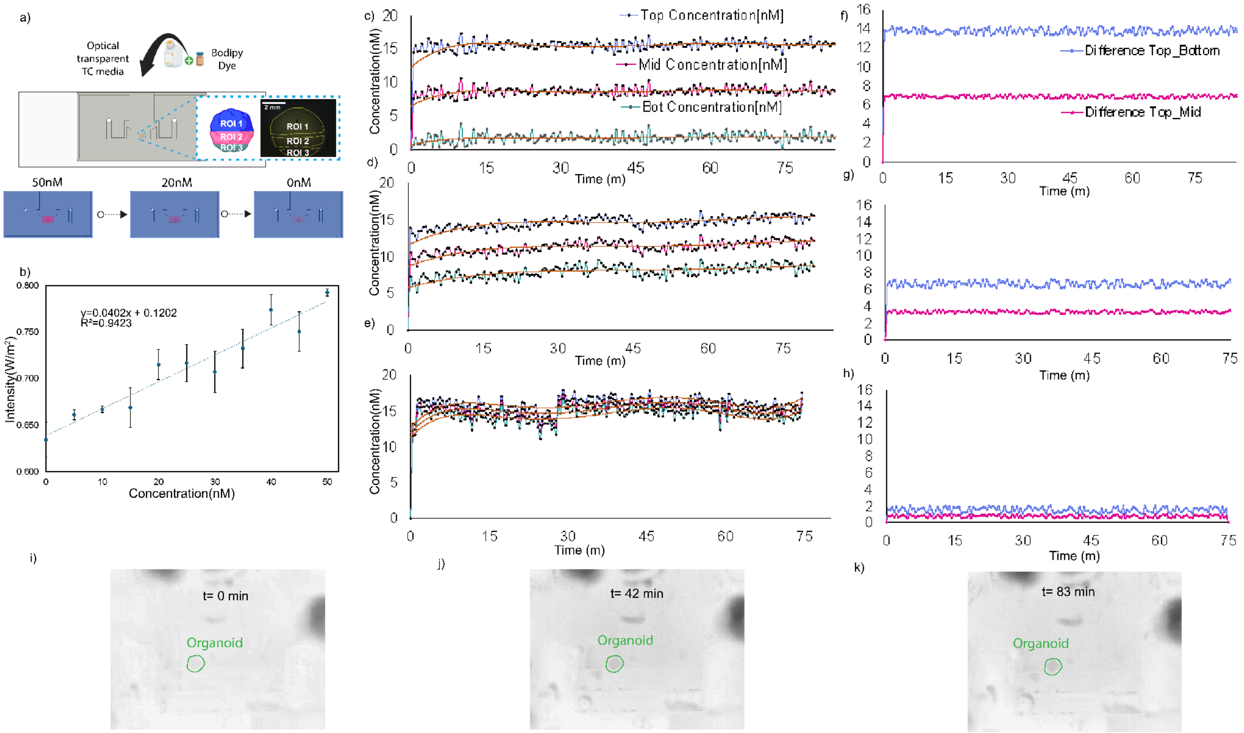


**Supplementary Figure 7. Quantification of BODIPY dye diffusion and concentration dynamics within the microfluidic chip**. (a) Schematic of the experimental setup showing the optical transparent tissue-culture (TC) medium channel loaded with BODIPY dye at three concentrations (50 nM, 20 nM, and 0 nM) and three defined regions of interest (ROI 1-3) within the observation area. Fluorescence images show the distribution of the BODIPY dye across the ROIs. (b) Calibration curve correlating fluorescence intensity (W/m$^2$) with dye concentration (nM), showing a linear relationship ($R^2$ = 0.9423) used to convert image intensity into absolute concentration values. (c) Temporal evolution of BODIPY dye concentrations at different depths within the channel—top, middle, and bottom layers—demonstrating the gradual stabilization of concentration profiles over the course of 75 minutes for initial concentration, (f) with preloaded concentration after 90 minutes, and with (g) preloaded concentration after 180 minutes. (f-h) Concentration difference between the top, middle, and bottom regions, showing diffusion-driven homogenization of the dye over time and minimal residual concentration gradients at later time points. (i-k) Bright-field images showing a representative organoid within the channel at 0, 42, and 83 minutes, highlighting the spatial stability of the sample during dye diffusion. Concentration values around the organoid increase gradually as equilibrium is reached, confirming that diffusion dominates over convective transport under static conditions.

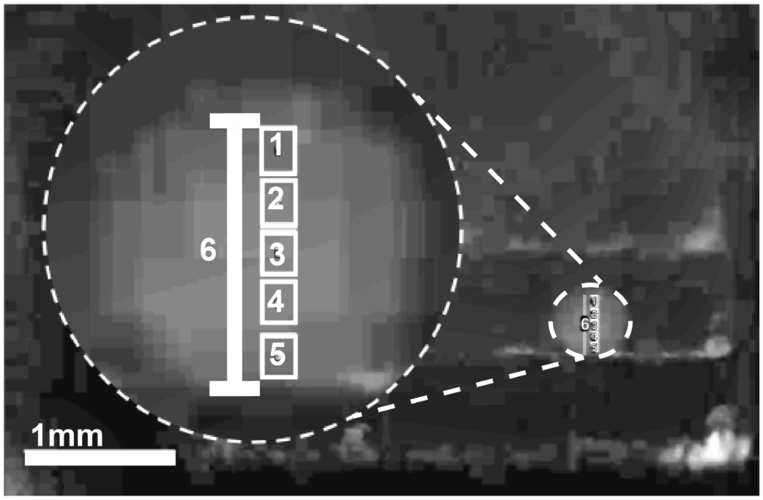


| | Pixel Intensity | Length |
|---|---|---|
| **1** | 35.38462 | 0.34376 |
| **2** | 41.67424 | 0.25782 |
| **3** | 42.52273 | 0.17188 |
| **4** | 40.93939 | 0.08594 |
| **5** | 35.97727 | 0 |

**Supplementary Figure 8. Calibration and spatial distribution of pixel intensity across the organoid cross-section.** A representative fluorescence image shows the spatial profile of Bodipy dye intensity across a single organoid (dashed circle). Five equally spaced regions of interest (ROIs 1-5) were drawn along the organoid's vertical axis to quantify the diffusion gradient from the periphery (ROI 1) toward the center (ROI 5). The table summarizes the mean pixel intensity and normalized distance (Length) for each ROI, where a gradual decrease in intensity toward ROI 5 indicates reduced dye penetration and concentration in the organoid core. The intensity profile shows a near-symmetric distribution with higher fluorescence at intermediate depths (ROIs 2-3, \~ 41-42 a.u.), suggesting partial equilibration of BODIPY diffusion. In comparison, peripheral (ROI 1) and central (ROI 5) regions exhibited lower fluorescence (\~ 35 a.u.). Scale bar = 1 mm.

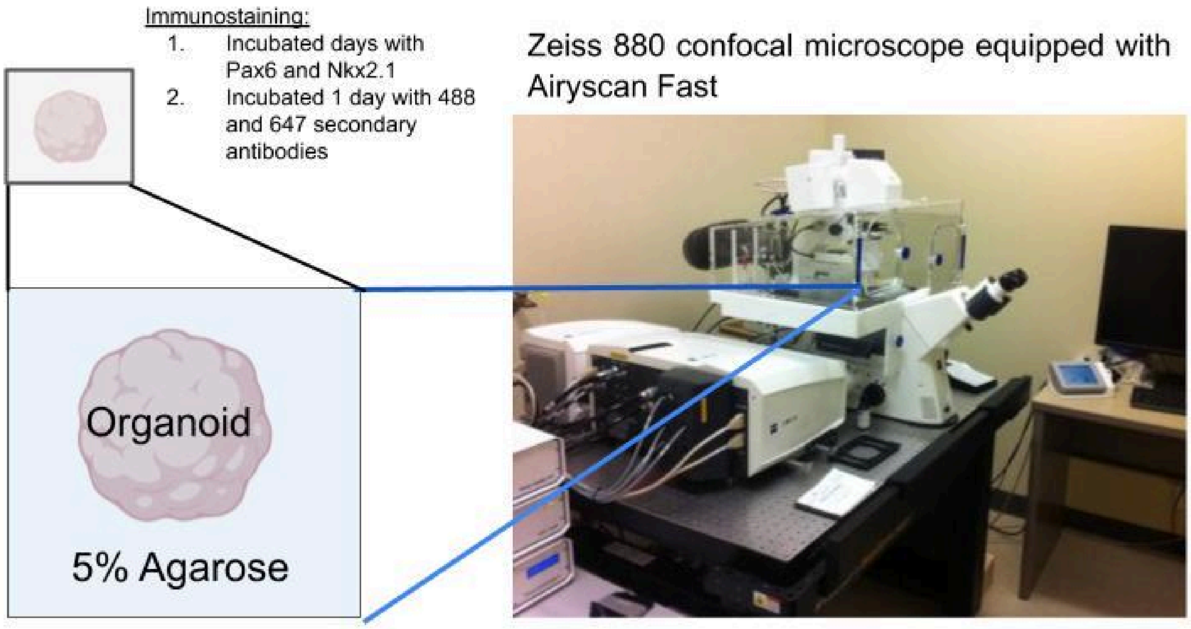


**Supplementary Figure 9. Workflow for organoid immunostaining and high-resolution imaging.** Organoids were embedded in 5% agarose and processed for immunostaining by incubating with primary antibodies against PAX6 and NKX2.1 for multiple days, followed by a 1-day incubation with Alexa Fluor 488- and 647-conjugated secondary antibodies. After staining, organoids were imaged using a Zeiss LSM 880 confocal microscope equipped with Airyscan Fast, enabling high-resolution three-dimensional reconstruction of marker distribution.
